# Transcriptomic signatures of Aβ- and tau-induced neuronal dysfunction reveal inflammatory processes at the core of Alzheimer’s disease pathophysiology

**DOI:** 10.1101/2023.09.15.557737

**Authors:** Lazaro M. Sanchez-Rodriguez, Ahmed F. Khan, Quadri Adewale, Gleb Bezgin, Joseph Therriault, Jaime Fernandez-Arias, Stijn Servaes, Nesrine Rahmouni, Cécile Tissot, Jenna Stevenson, Hongxiu Jiang, Xiaoqian Chai, Felix Carbonell, Pedro Rosa-Neto, Yasser Iturria-Medina

## Abstract

Molecular mechanisms enabling pathology-induced neuronal dysfunction in Alzheimer’s disease (AD) remain elusive. Here, we use mechanistic computational models to infer the combined influence of PET-measured Aβ and tau burdens on fMRI-derived neuronal activity and to subsequently identify the transcriptomic spatial correlates of AD pathophysiology. Our results reveal overrepresented genes and biological processes that participate in synaptic degeneration and interact with Aβ and tau deposits. Furthermore, we confirmed the central role of the immune system and neuroinflammatory pathways within AD pathogenesis; microglia were significantly enriched in the gene set associated with Aβ and tau synergistic influences on neuronal activity. Lastly, our computational approach unveiled drug candidates with the potential to halt or reduce the observed pathological effects on neuronal activity, including existing medication for cancer, immune disorders, and cardiovascular diseases, many currently under clinical evaluation in AD. Overall, these findings support the notion that the AD brain experiences functional changes intricately associated with a diverse spectrum of molecular processes.

## Introduction

Alzheimer’s disease (AD) seems to be determined by multiple interacting molecular elements and generalized dysfunction (Calabrò et al., 2021; Iturria-Medina et al., 2022). Notably, 91% of all molecular pathways recorded in the Kyoto Encyclopedia of Genes and Genomes (KEGG) have been found to be associated with AD in at least 5 studies (Morgan et al., 2022). In cell culture, animal and post-mortem research, mechanisms of neuroinflammation/immune system, metabolism, cholinergic synapse, cancer, diabetes and chemokine signaling typically rank high in associations with AD (Calabrò et al., 2021; Morgan et al., 2022; Santiago & Potashkin, 2021). However, these results may not fully capture disease progression in the living human organism.

Neuronal dysfunction in AD is associated with toxic protein accumulation, including amyloid beta (Aβ) plaques and tau neurofibrillary tangles (NFTs) (Jack et al., 2018; Maestú et al., 2021). PET-measured Aβ and tau aggregation in the brain serve to predict cognitive decline (Chandra et al., 2019). *In-vivo* animal experiments and modeling approaches suggest that Aβ and tau also synergistically interact to impair neuronal (Maestú et al., 2021; Targa Dias Anastacio et al., 2022; van Nifterick et al., 2022), with Aβ and tau pathologies likely prompting brain network hyperactivity as the disease progresses (Busche & Hyman, 2020; Tok et al., 2022; Vossel et al., 2017). Although these effects are consistent across the literature, limitations to concurrently measure neuronal activity alterations, pathological severity, and molecular profiles in the living human brain represent a major obstacle towards clarifying AD mechanisms (Gabitto et al., n.d.; Iturria-Medina et al., 2022; Maestú et al., 2021; Nandi et al., 2022). Lacking definitive pathways to target, the incomplete characterization of AD’s pathophysiology may have contributed to the limited efficacy of some of the thus-far proposed therapeutics (Cummings et al., 2021; Iturria-Medina et al., 2018). Increasing the understanding of the affected biological processes will also allow their early modification through healthy lifestyle choices and clinical monitoring, boosting disease prevention (Silva et al., 2019; *World Alzheimer Report 2022 – Life after Diagnosis: Navigating Treatment, Care and Support*, n.d.).

Integrative computational modeling of *in-vivo* human pathophysiological processes offers a powerful alternative to circumvent experimental shortcomings in AD research (Adewale et al., 2021; Carbonell et al., 2018; Deco et al., 2018; Iturria-Medina et al., 2021, 2022; Khan et al., 2022; Lenglos et al., 2022; Sanchez-Rodriguez et al., 2018; Sotero & Trujillo-Barreto, 2008; Stefanovski et al., 2019). We recently used personalized computational models to decode synergistic Aβ and tau effects on neuronal excitability in AD progression (Sanchez-Rodriguez et al., 2023). This allowed us to robustly infer *in-vivo* patient-specific values of neuronal excitability and describe their associations with pathological severity, disease biomarkers (e.g., p-tau217, p-tau231) (Zetterberg & Blennow, 2021) and altered electroencephalographic indexes (Babiloni et al., 2013; Sanchez-Rodriguez et al., 2018). The obtained Aβ and tau weights also predicted cognitive decline in the AD-related cohort. However, the precise molecular pathways by which AD pathology impacts neuronal excitability throughout the brain remain largely uncharacterized.

In this study, we aimed to describe the biological underpinnings of neuronal activity alterations in AD’s pathophysiology. First, we utilize generative brain models to estimate the combined spatiotemporal influence of Aβ and tau (measured via PET) on neuronal activity (measured through fMRI biomarkers) for cognitively unimpaired and AD participants. Second, we use whole-brain transcriptomics to identify genes with spatial expressions predicting the regional neuronal activity effects of Aβ, tau and their synergistic interaction, respectively. We examine this evidence in the context of the biological mechanisms that may be associated with AD’s development. This analysis results in a clear and consistent Aβ+tau → neuronal-activity molecular profile, with both distinctive mechanisms and processes shared with diseases such as infection, cancer and retinal conditions. Major associations with immune system, cell communication and developmental mechanisms exist, driven by the synergistic interaction of Aβ and tau. Third, we detect the cell types that are most likely related to neuronal activity alterations by the causal combined roles of Aβ and tau pathologies, observing a predominant role of microglia. Fourth, we identify potential pharmacological interventions repurposing existing drugs to modify the diseased biological processes. Collectively, our computational experiments demonstrate the complexity of the disease and characterize its diverse biological affectations profile. This comprehensive computational approach discovers fundamental *in-vivo* disease mechanics to target with advanced therapeutics.

## Results

### Neuronal activity alteration patterns in AD reveal molecular disease signatures

We obtained *in-vivo* structural and functional MRI, Aβ and tau-PET and clinical evaluations for 47 cognitively unimpaired (CU) and 16 AD participants (Supplementary File 1—table 1) from the Translational Biomarkers in Aging and Dementia cohort (TRIAD, https://triad.tnl-mcgill.com/). In addition, we processed bulk transcriptomic data for the whole adult human brain from the Allen Human Brain Atlas (AHBA) (Adewale et al., 2021; Allen Human Brain Atlas, 2013). We used a personalized computational model to decode the causal spatial influence of Aβ, tau and their combined synergistic interaction on neuronal activity (modeled as the regional multiplication of the Aβ and tau burdens, Aβ·tau) (Sanchez-Rodriguez et al., 2023) –see Figure 1. For each AD and CU subject in the TRIAD database, we assumed that neuronal excitability across the brain regions was potentially influenced by the local Aβ and tau accumulations. These alterations spatiotemporally transmit through intra-regional and cortico-cortical connections (derived from diffusion MRI). The in silico pathophysiological excitatory and inhibitory activities (Wilson & Cowan, 1972) were transformed into blood-oxygen-level-dependent (BOLD) signals (Sotero & Trujillo-Barreto, 2007; Valdes-Sosa et al., 2009) and fitted to the subject’s real BOLD signal content in the physiologically-relevant neuronal activity range (0.01–0.08 Hz) (Yang et al., 2018). Subsequently, we identified distinctive spatial Aβ, tau and Aβ·tau neuronal activity alterations patterns via statistical evaluation of their induced neuronal excitability perturbations in the AD vs CU groups.

**Figure 1.**
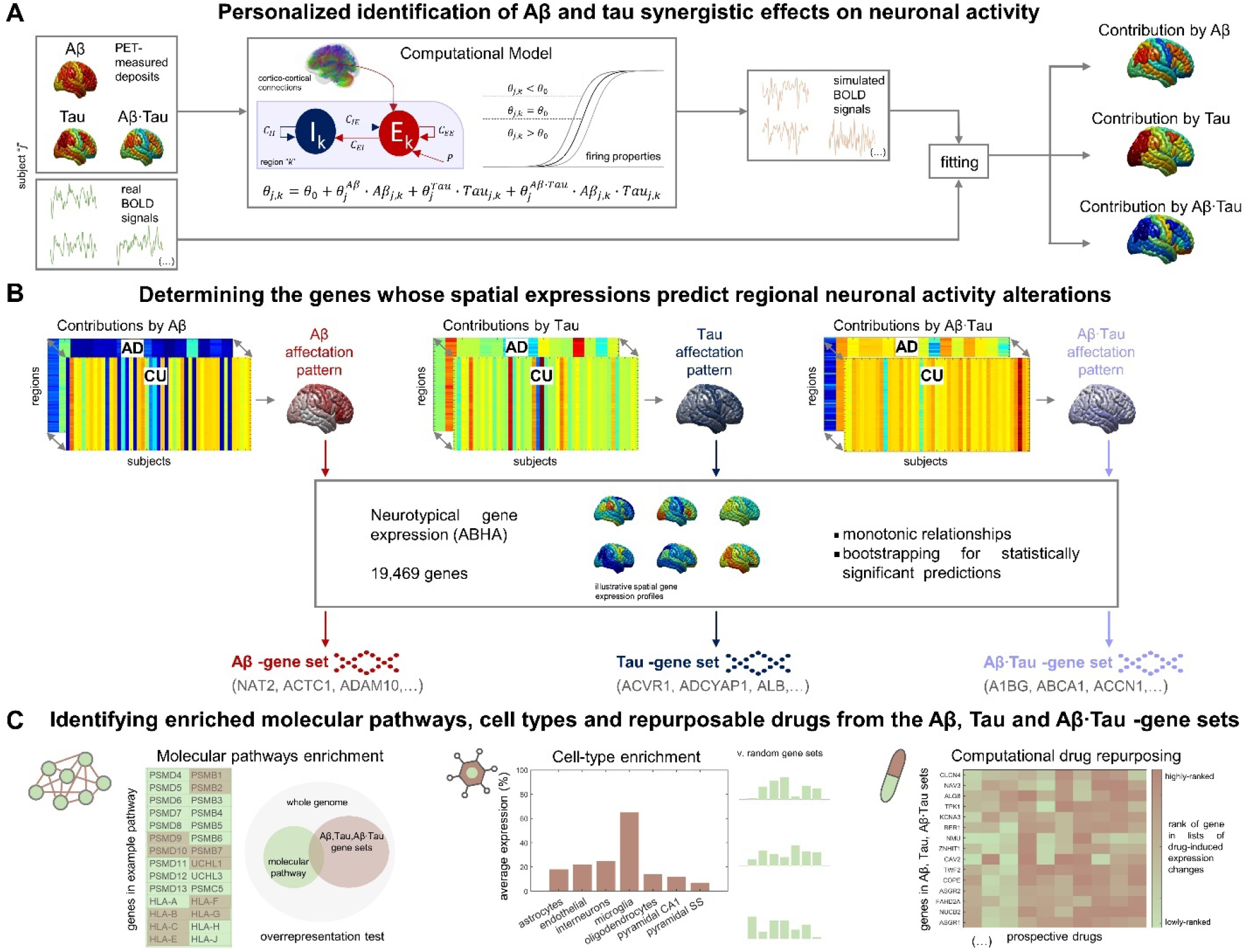
Methodological approach for determining the molecular mechanisms associated with Aβ and tau-induced neuronal dysfunction in AD. (**A**) For each participant, neuronal excitability within a brain region depends on the combined Aβ and tau accumulations. Simulated excitatory and inhibitory activities are transformed into BOLD signals. The most-likely *in-vivo* subject-specific Aβ and tau effects are obtained through maximizing the similarity with the participant’s real fMRI across regions. (**B**) Statistical comparison of the obtained regional Aβ, tau and Aβ·tau contributions to pathophysiological neuronal activity between the AD and CU groups yields spatial alterations patterns by each of these disease factors (the higher the statistic, the more different the groups are). Next, we investigate spatial correlations with neurotypical whole-brain transcriptomics (99% bootstrap confidence intervals) and obtain the genes which expressions predict the regional neuronal activity effects by Aβ, tau and Aβ·tau. (**C**) The sets of Aβ, tau and Aβ·tau molecular associates serve to study enriched biological processes (molecular pathways from multiple gene ontologies being overrepresented in these gene lists) (Zhou et al., 2019), brain cell-types (the Aβ, tau and Aβ·tau gene sets having higher expression for a particular cell type than what is expected by chance) (Skene & Grant, 2016) and prospective repositioned pharmacological agents to halt or reduce AD-affected processes (by comparing the gene sets to databases of ranked gene lists for drug-induced gene expression changes) (Evangelista et al., 2022).

We aimed to determine the genes from the human transcriptome whose spatial expressions predict the regional neuronal activity effects by each pathophysiological factor. For this purpose, we computed 99% bootstrap confidence intervals (99CI) for the brain-wide correlations between the Aβ, tau and Aβ·tau spatial patterns and the expression of each gene in the AHBA transcriptome. We identified 756, 650 and 1987 genes, respectively, in the Aβ, tau and Aβ·tau-associated gene sets (99CI did not include zero). The lists, provided in Supplementary File 2, include several genes that might affect AD risk (Calabrò et al., 2021). For instance, the microglial activation modulator *CD33* (Sialic Acid-Binding Ig-Like Lectin 3) is one of the top-ranked genetic factors identified in AD genome-wide association studies (Zhao, 2019). Gene *ADAM10* (α disintegrin and metalloproteinase domain-containing protein 10) plays a critical role in cleavage of the amyloid precursor protein (APP) (Calabrò et al., 2021). *SNCA* (synuclein Α) is essential for presynaptic signaling and membrane transport and participates in NFT formation and Aβ deposits (Calabrò et al., 2021). Finally, the protein encoded by the gene *CLU* (clusterin) inhibits Aβ fibrils formation (Calabrò et al., 2021). As reported in the next subsections, the Aβ, tau and Aβ·tau molecular associates of AD pathogenesis were further investigated in terms of overrepresented biological mechanisms, cellular types associated with brain-wide functional affectations and repositioned drug candidates with potential therapeutic benefit.

### Immune and cell communication processes relate to AD pathology-induced neuronal dysfunction

Despite the moderate isolated influence of single molecules, multifactorial and complex disorders like AD are more recently approached in terms of comprehensive biological processes expressing the disease’s signatures (Calabrò et al., 2021; Iturria-Medina et al., 2022; Morgan et al., 2022). We proceeded to compare the three neuronal activity alterations-associated gene sets to ontology terms from various sources in Metascape (Zhou et al., 2019), detecting the molecular pathways that are significantly overrepresented in the combination of such genetic signatures (*Methods, Statistical Analyses,* Supplementary File 3). The top 20 enriched functional clusters that were retrieved, together with the gene lists where the pathways were found statistically significant (hypergeometric tests, FDR-corrected, q< 0.05) are shown in Figure 2A. Supplementary File 1—table 2 presents the Aβ+tau → neuronal-activity genes that are consistently involved (95% percentile) within the top statistically significant identified biological pathways.

**Figure 2.**
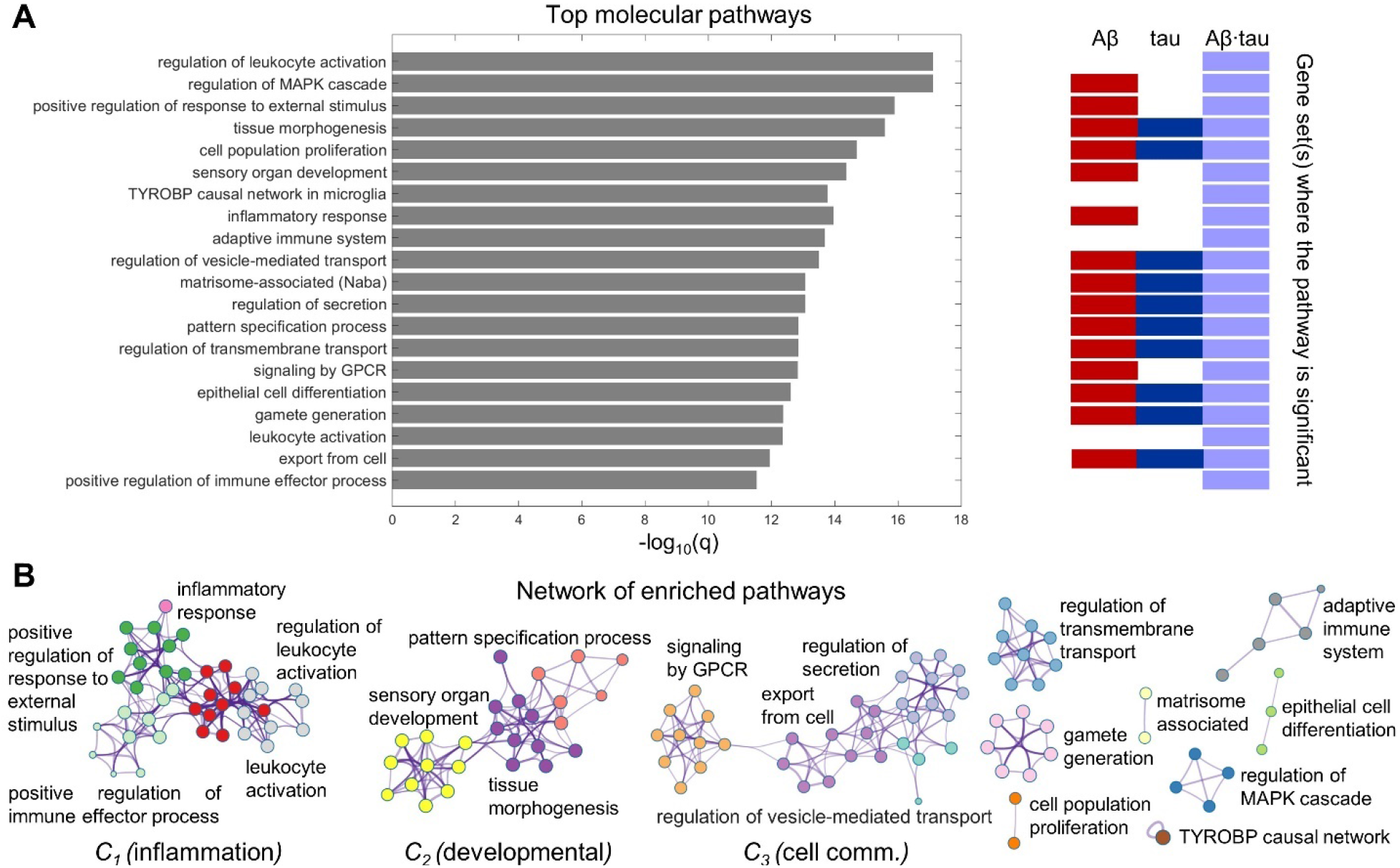
Neuroinflammation pathways emerge as major processes associated with Aβ-tau interactions. (**A**) Top 20 pathways clusters from multiple gene ontologies that are enriched in the combined Aβ-, tau- and Aβ·tau-associated gene sets (hypergeometric tests, q < 0.05, Benjamini-Hochberg corrected). The representative biological processes (term with the lowest *p*-value within a cluster) are used as labels. Additionally, the specific gene set(s) for which the pathways are statistically significant have been indicated next to the bar graph. (**B**) Network plot showing the intra-cluster and inter-cluster similarities among the obtained molecular processes. Each node represents an enriched pathway. The network is colored by the cluster labels, which are written next to each cluster. Major clusters include neuroinflammation and immune system processes (C_1_), developmental pathways (C_2_) and cell communication mechanisms (C_3_).

Biological processes with a high genetic overlap are also visualized in a connected network (Figure 2B) reflecting the relatedness of separate pathways (Zhou et al., 2019). In this manner, we observe the clustering of various neuroinflammation and immune system pathways, i.e., *inflammatory response* biological processes connect with the *positive regulation of immune effector process*, *positive regulation of response to external stimulus, leukocyte activation* (and its *regulation*) pathways, given the high similarity of their enriched terms. Persistent chronic inflammation, due to genetic and lifestyle factors, plays a key role at the onset and later progression of neurodegeneration (Calabrò et al., 2021; Newcombe et al., 2018). Notably, Aβ and tau accumulation can both trigger and be triggered by disbalanced inflammatory signals (Newcombe et al., 2018). Another functional cluster consists of developmental processes (*sensory organ development*, *tissue morphogenesis*, *pattern specification process*). Cell communication/transport mechanisms that are fundamental to proper synaptic function and are implicated in AD pathogenesis according to several reports (Gadhave et al., 2021) were also pinpointed among the top enriched molecular processes in a major cluster (*regulation of secretion, regulation of vesicle-mediated transport, export from cell, signaling by GPCR*). Figure 2 summarizes the comprehensive view of the molecular mechanisms that associate with the causal combined roles of Aβ and tau pathologies on AD’s neuronal activity alterations.

Additionally, we examined biological processes separately related to the Aβ, tau and Aβ·tau gene sets. These results appear in Supplementary Files 4-6. Immune system pathways were once again overrepresented in the Aβ·tau set, while developmental and synaptic processes were enriched for Aβ’s molecular associates. Notably, some pathways that ranked lower in the integrative analysis in Figure 2, had relevant associations with the tau-associated gene list (with less elements than the Aβ and Aβ·tau molecular signatures). Amongst the enriched terms, several supposedly tau-related processes (R. E. Bennett et al., 2018; Mandelkow & Mandelkow, 2011) including *cortical cytoskeleton organization*, *regulation of actin filament organization*, *blood vessel development* and *post-translational protein phosphorylation* appeared.

We also explored molecular associations with other diseases according to the genes predicting the spatial neuronal activity combined Aβ and tau effects. We determined which disease pathways, curated in DisGeNET (Piñero et al., 2017; Zhou et al., 2019), were enriched in our gene sets (Supplementary File 1—figure 1, Supplementary Files 7-9). Notably, the obtained enriched terms include several infection and immunological conditions (e.g., immunosuppression, Behcet syndrome and lupus), certain cancers, and eye diseases, for the three considered sets of molecular associates. Likewise, we retrieved emblematic AD phenotypical symptoms (D. A. Bennett et al., 2013; Ghiso & Frangione, 2002) like *memory impairment* (enriched in both Aβ and tau signatures) and *amyloidosis* (Aβ), further demonstrating that our approach unifying whole-brain transcriptomics, molecular and functional neuroimaging, and personalized computer-simulated neuronal activity reliably identifies affected biological mechanisms.

### Pyramidal cells, interneurons and microglia are vulnerable to the Aβ, tau and Aβ·tau molecular associates

Next, we hypothesized that the gene sets associated with each of the pathophysiological neuronal activity patterns would be particularly enriched in distinct cell types. We performed a bootstrapping-based cell type enrichment analysis on the Expression Weighted Celltype Enrichment toolbox (Skene & Grant, 2016) and determined the statistical likelihood of brain cell types being enriched compared to a background gene set (transcriptome obtained from the somatosensory cortex and hippocampus CA1) (Figure 3).

**Figure 3.**
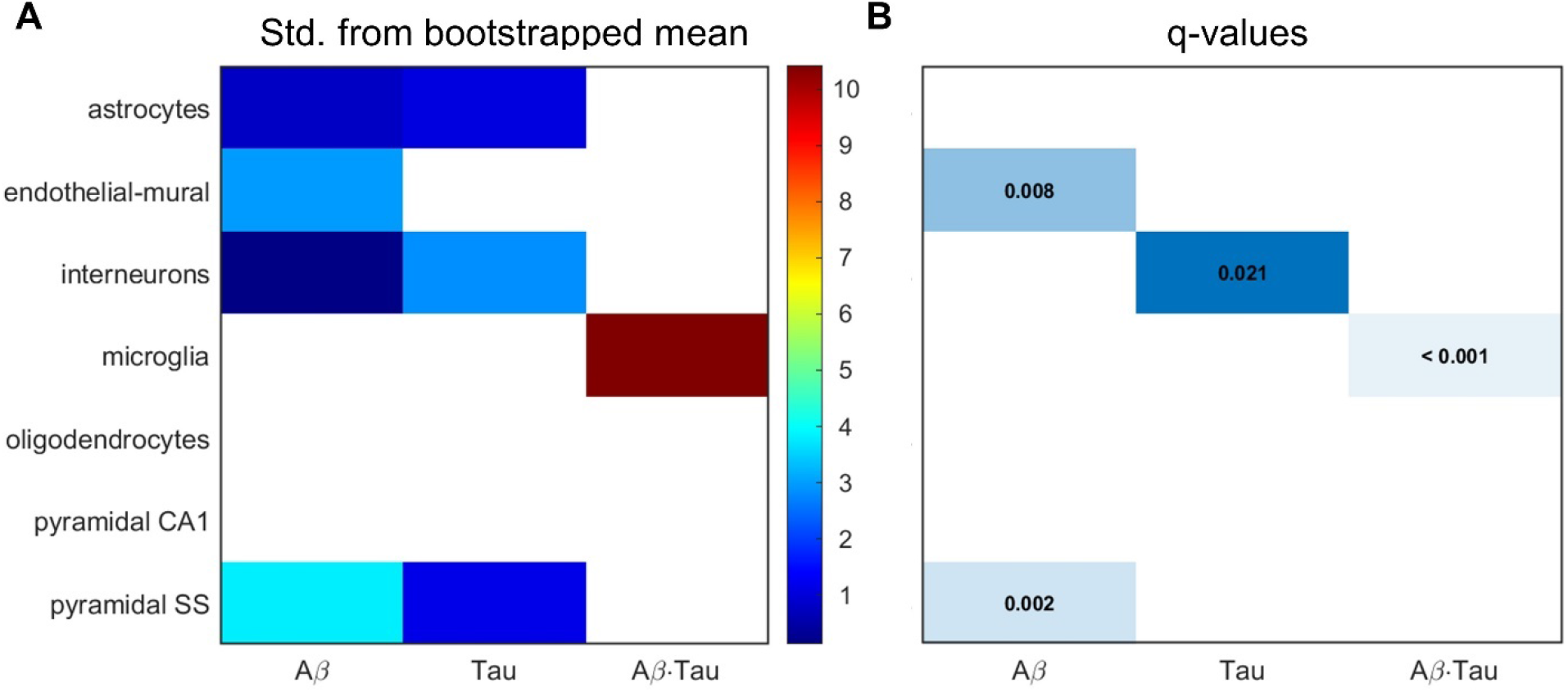
Transcriptomic associates of Aβ- and tau-induced neuronal activity alterations converge to neuro-vascular cellular compartments. Results of bootstrapping tests to evaluate the probability of the Aβ-, tau- and Aβ·tau-associated gene sets having higher expression for a particular brain cell type than what is expected by chance. (**A**) Number of standard deviations from the bootstrapped mean for every gene set and cell type. Non-white boxes indicate that the given molecular associates’ expression in the specific cell type is, on average, higher than that of the bootstrapped sets. (**B**) Statistically significant enrichment (q < 0.05, Benjamini-Hochberg corrected).

We found strong evidence supporting pyramidal cells (q = 0.002 and δ = 3.804, denotes the number of standard deviations which the average expression of the gene list falls from the bootstrapped mean) and endothelial-mural cells, the constituent of blood vessels (q = 0.008 and δ = 2.950), as the cell types most enriched amongst the Aβ molecular associates. Previous experiments (Koizumi et al., 2016) show impairment to cerebral blood vessels by extracellular buildup of Aβ, while vascular dysfunction may promote Aβ accumulation in a detrimental feedback loop. Pyramidal neurons, the most abundant neural cells in the cortex, are known to be a preferential target for both Aβ and tau toxic deposits (Maestú et al., 2021). In consequence, we would expect statistically significant overrepresentation in both cases. However, only interneurons presented significant enrichment for the tau susceptibility genes, according to the bootstrapping analysis (q = 0.021 and δ = 2.834). Phosphorylated tau seems to accumulate early in hippocampal interneurons of AD patients, impairing adult neurogenesis and circuital function (Xu et al., 2020; Zheng et al., 2020). On the other hand, the Aβ·tau molecular signature had significant microglial expression (q < 0.001 and δ = 10.425). To our knowledge, functional Aβ and tau interactions have never been studied in the context of genetic cell enrichment although analyses of the disease’s polygenic post-mortem expression have also found damage to microglia (Galatro et al., 2017; Newcombe et al., 2018).

Our observations will necessitate further validation to fully comprehend the causal cell-specific synergistic effects of Aβ and tau. For this purpose, a brain-wide multimodal cell type atlas of AD could be available from The Seattle Alzheimer’s Disease Cell Atlas (SEA-AD) consortium in the future (Gabitto et al., n.d.). Different pathophysiological mechanisms may determine specific cellular vulnerability patterns.

### Repurposed immunologic drugs could halt or reduce AD

Finally, we examined whether existing drugs could be repurposed to temper AD’s observed pathological effects. For this purpose, we computationally searched for chemical compounds with well-described mechanisms of action maximally up or down regulating the expression of all the genes in the Aβ+tau → neuronal-activity molecular profile, utilizing the webserver SigCom LINCS (Evangelista et al., 2022). In Figure 4, we report the top statistically significant (q < 0.05) candidate drugs for LOAD that have been FDA-approved for intended use elsewhere (accessed through https://pubchem.ncbi.nlm.nih.gov/ on May 10^th^, 2023). The full list of compounds retrieved from SigCom LINCS and additional details as doses and other experimental conditions for drug mechanisms characterization can be found in Supplementary Files 10-11, while drug indications and blood-brain barrier (BBB) permeabilities (Meng et al., 2021) of the top prospective medications are provided in Supplementary File 12. In separate analyses, we also queried drug-molecular targets interactions of the independent Aβ, tau and Aβ·tau-associated gene sets (Supplementary Files 13-18).

**Figure 4.**
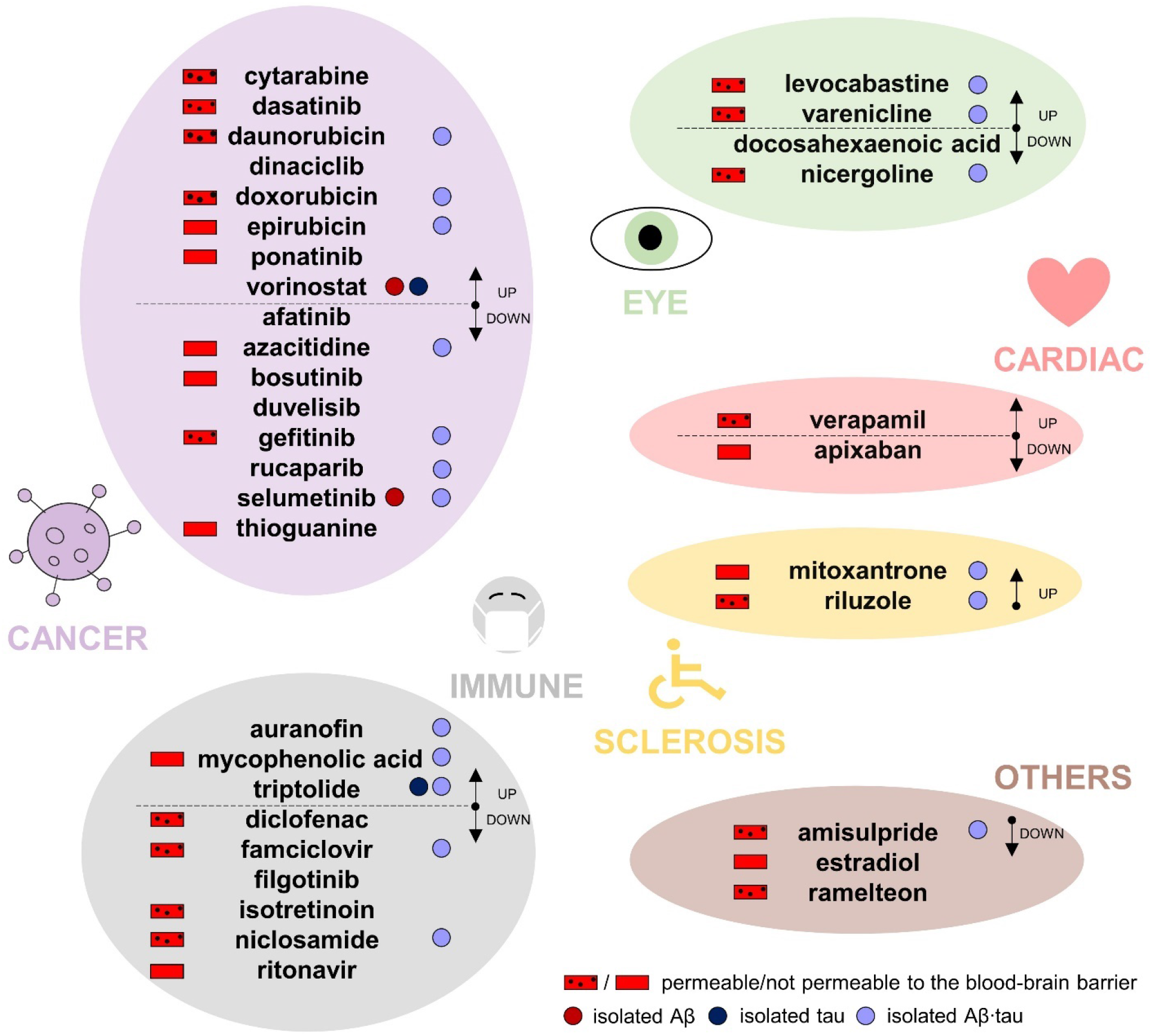
Top drug repositioning candidates to overcome the observed Aβ+tau → neuronal-activity affectation profile. Reported are existing drugs which molecular interactions would induce gene expression changes in the set of all Aβ-, tau- and Aβ·tau-associated genes (Mann– Whitney U test, q < 0.05, Benjamini-Hochberg corrected). The predicted chemical compounds have been organized in major groups according to their drug use indications. All medications are FDA-cleared for either treatment of cancer, various immune system/infection/inflammatory processes (“immune”), eye diseases, cardiovascular conditions, multiple or amyotrophic lateral sclerosis or “other” disorders. The groups are further divided by whether the candidate drug up- or down-regulates the genes linked to the neuronal activity alterations by AD. Additionally, blood-brain barrier permeability, when this information was available, is specified next to the name of the drug (red rectangles on the left). Drugs that could also target the separate Aβ, tau or Aβ·tau molecular associates are identified with accordingly colored circles on the right of the compound’s name, e.g., the chemical selumetinib may be used to attack Aβ- and Aβ·tau-associated gene sets.

The chemical compounds interacting with LOAD’s spatial molecular associates and potentially inducing therapeutic changes are, mostly, drugs used for the treatment of cancer and immune system-related disorders. Common indications among these medications include leukemia, lymphoma and breast cancer. In clinical research, prospective disease-modifying AD drugs commonly target cancer pathways (Morgan et al., 2022). Computational drug repurposing studies have similarly assessed the benefits of anti-cancer drugs. For example, a multi-omics study identified interactions of afatinib, dasatinib, gefitinib and ponatinib with AD-affected genes (e.g., *APP*, *SNCA*) (Advani & Kumar, 2021). Among the top immunological drug candidates, the immunosuppressant medication mycophenolic acid, indicated for prophylaxis of organ rejection, has been reported to attenuate neuronal cell death (Ebrahimi et al., 2012); diclofenac could potentially associate with reduced AD risk and slower cognitive deterioration (Rivers-Auty et al., 2020), while antiherpetic medication as famciclovir may also prevent AD incidence (Calabrò et al., 2021; Linard et al., 2022). Likewise, anti-inflammatory multiple sclerosis medication has shown promise in AD mouse models for reversing all Aβ, tau and microglia pathologies, and synaptic and cognitive dysfunction (Dionisio-Santos et al., 2021; Leßmann et al., 2023). It is worth noticing that, in clinical trials, drugs with anti-inflammatories properties have not slowed cognitive and/or functional decline (Howard et al., 2020; Melchiorri et al., 2023). One possible explanation is that the thus-far tested agents interfere with microglia’s supportive function instead of modulating its detrimental chronic activation effects (Howard et al., 2020; Melchiorri et al., 2023; Rivers-Auty et al., 2020; Shen et al., 2018). At least 18 investigational drugs targeting neuroinflammation currently undergo clinical assessment, including phase III trials (Melchiorri et al., 2023; Reading et al., 2021).

Among the resting identified prospective candidates, cardiovascular drugs may lower the incidence of dementia –apixaban (Bezabhe et al., 2022)– and delay progression in a mouse model of AD –verapimil (Ahmed et al., 2021). Additionally, docosahexaenoic acid (omega-3) supplementation has been linked to reduced AD risks (Arellanes et al., 2020; Quinn et al., 2010). Randomized trials finding interactions with *APOE4* suggest that such AD carriers could also potentially present favorable imaging and cognitive outcomes with high dose docosahexaenoic acid supplementation treatments (Arellanes et al., 2020). On the other hand, retinopathy, glaucoma and age-related macular degeneration are deemed prominent signs of AD pathology (Mirzaei et al., 2020), functionally sharing affected molecular pathways (Supplementary File 1—figure 1, Supplementary Files 7-9), which explains the appearance of visual impairments medication among the top prospective drugs. Overall, our results signal possible therapeutic strategies to be tested in randomized controlled trials (RCTs) for the treatment and prevention of AD, bypassing the early stages of drug design for compounds with known pharmacokinetic/pharmacodynamic properties.

## Discussion

A cardinal problem of modern neuroscience is understanding brain disease to a degree that enables effective therapeutic interventions. In AD, research has suggested a myriad of interacting mechanisms with potential central contributions by Aβ and tau pathological deposits (Iturria-Medina et al., 2018; Maestú et al., 2021; Newcombe et al., 2018; Sanchez-Rodriguez et al., 2023; Therriault et al., 2022). We investigated combined neuronal activity alterations by Aβ and tau pathologies in an AD patient cohort and mapped their relationship to molecular processes through an integrative computational approach informed by *in-vivo* neuroimaging and whole-brain transcriptomics. The estimation of the Aβ+tau → neuronal-activity molecular affectations profile provides privileged insights into *in-vivo* human AD pathophysiology.

Several of the identified spatial molecular correlates from the human brain transcriptome have been linked to AD in the past (Calabrò et al., 2021). For example, the *SNCA* gene translates into the presynaptic protein α-synuclein, which presents high concentration in the cerebrospinal fluid of mild cognitive impairment and AD patients and forms deposits that have been found in the majority of autopsied AD brains (Twohig & Nielsen, 2019). Another possible AD genetic modifier, clusterin, seems to enhance excitatory synaptic transmission while reducing Aβ pathology (F. Chen et al., 2021). Gene *RIPK2* is a mediator of mitochondrial dysfunction in oligodendrocytes and demyelination (Natarajan et al., 2013), *SYK* coordinates neuroprotective microglial response to Aβ pathology (Ennerfelt et al., 2022) and *ANXA1* plays an important role in controlling neuronal damage by immune responses (You et al., 2021). All these molecules are central to the top overrepresented biological processes (Supplementary File 1—table 2) and the overall AD *in-vivo* molecular signature.

Collectively, the enriched pathways within AD’s Aβ+tau → neuronal-activity molecular profile might represent the biological processes that are most likely associated to diseased brain function. We utilized the most comprehensive and robust resources for the analysis of the discovered molecular profile (Evangelista et al., 2022; Skene & Grant, 2016; Zhou et al., 2019) –e.g., Metascape integrates major current biological databases including KEGG Pathway, GO Biological Processes, WikiPathways and PANTHER Pathway. While our findings rely on the consulted databases, we observed congruence across analyses and with the existing literature. We have confirmed existing hypotheses regarding AD as a generalized condition (Calabrò et al., 2021; Morgan et al., 2022), although certain types of molecular pathways seem to play a fundamental role in AD’s pathophysiology. Memory impairment and other hallmark AD signs including amyloidosis and phosphorylation (D. A. Bennett et al., 2013; Ghiso & Frangione, 2002; Mandelkow & Mandelkow, 2011), were overrepresented in Aβ and tau’s separate molecular associates of pathophysiological neuronal activity. Most crucially, neuronal activity alterations by Aβ, tau and their synergistic interaction were consistently related to inflammation processes.

In effect, the spatial molecular associates of the interaction between Aβ and tau pathologies were more enriched for microglial expression than expected by chance. When activated, microglia may trigger different processes that play a role in AD risk (Calabrò et al., 2021; Kwon & Koh, 2020; Shen et al., 2018). The M2 activated phenotype is believed to protect the brain versus chronic neuroinflammation. The M1 phenotype, on the other hand, increases the secretion of pro-inflammatory cytokines. Studies have suggested that prolonged, uncontrolled immune responses cascade to modify physiological properties and the neuronal activity balance through interactions with Aβ and tau (Calabrò et al., 2021; Kwon & Koh, 2020; Newcombe et al., 2018; Shen et al., 2018). Our findings indicate that neuroinflammation also interplays with Aβ and tau synergistic effects, which seems to be a key factor in AD’s pathophysiology (Busche & Hyman, 2020; Sanchez-Rodriguez et al., 2023). The identification of a major cluster of immunological pathways within AD’s neuronal activity molecular signatures warrants further investigation. In our previous work (Sanchez-Rodriguez et al., 2023), we sought to decode possible neuroinflammatory influences (interacting with Aβ and tau effects) to neuronal activity through personalized computational models. However, only slight significant differences in the translocator protein microglial activation -PET data existed between AD and CU subjects, underscoring broadly discussed limitations of PET tracers being unspecific to inflammatory variants as the protective M2 and the deleterious M1, which exacerbates the disease in late stages (Nutma et al., n.d.; Shen et al., 2018).

Further improvements and clinical validation are necessary for implementing treatment strategies stemming from computational modeling of neuropathological mechanisms (Iturria-Medina et al., 2018; Maestú et al., 2021). The dataset utilized in this study was collected at a specialist memory clinic that receives relatively young dementia patients. This highly specialized setting may pose a limitation in terms of generalizability, although subjects diagnosed as “early-onset” and/or “familial” AD were excluded from the current analysis. Additionally, the percentage of female subjects within the CU (AD) group was slightly higher (lower) than AD’s prevalence among women, i.e., nearly two-thirds of the total number of cases (*World Alzheimer Report 2022 – Life after Diagnosis: Navigating Treatment, Care and Support*, n.d.). Several factors contributed to this disproportion including the availability of volunteers and whether the necessary imaging modalities had been collected at the time of sample curation. Regarding the biophysical model for neuronal activity alterations due to AD’s pathology, we considered perturbations to pyramidal neurons only. Albeit a sound approximation given the pyramidal preponderance in the cortex (Maestú et al., 2021) –and with local connections propagating alterations to inhibitory populations as well (Wilson & Cowan, 1972)– this assumption could be relaxed by considering an inhibitory influence model and re-estimating the relevant pathophysiological parameters. By doing so, we may test hypotheses for inhibitory circuit impairment in AD (Maestú et al., 2021; Targa Dias Anastacio et al., 2022; Zheng et al., 2020). Importantly, we focused the scope of this investigation into AD mechanisms to influences by Aβ and tau only. The identified subject-specific Aβ and tau neuronal activity alterations should be interpreted as their causal pathophysiological effects disregarding other possible contributors, while the corresponding molecular associates characterize biological events that, in general, coexist with such alterations. It is known that additional factors as glial cell activity affect neuronal firing, even in healthy states (Targa Dias Anastacio et al., 2022). Our personalized models are readily modifiable (Sanchez-Rodriguez et al., 2023) to consider other pathological factors, provided that the corresponding brain imaging modalities are available. Advanced causal computational models unifying neuroimaging and omics exist (Adewale et al., 2021; Iturria-Medina et al., 2021, 2022; Khan et al., 2022; Lenglos et al., 2022), although they have yet to tackle the generation of (pathophysiological) neuronal activity. In future work, we intend to expand the high-dimensionality, multimodal approaches compiled within the in-house open-access NeuroPM-box software (Iturria-Medina et al., 2021) with quantification tools for unveiling molecular mechanics of pathological influences on neuronal activity.

The disease-oriented computational drug repurposing strategy that we present constitutes an accelerated alternative to costly drug development for AD as preliminary safety and bioavailability criteria are already established for the identified chemical compounds (Corbett et al., 2012; Mullen et al., 2016; Petralia et al., 2022). In 2021, approximately 40% of Alzheimer’s trials registered on ClinicalTrials.gov used repurposed existing medication (Cummings et al., 2021). Previous studies have assessed the usefulness of several of our discovered candidate pharmacological agents targeting affected AD pathways. Converging evidence indicates that cancer treatment may be related to a decreased risk of AD due to a pathophysiological overlap between both diseases, albeit a worsened cognition being in some studies linked to oncology drugs (D. Chen et al., 2021; Frain et al., 2017; Plun-Favreau et al., 2010). The FDA-approved compound dasatinib, for the treatment of chronic myeloid leukemia, has reduced tau pathology in mice (Roberts et al., 2021) and is the subject of an ongoing clinical study evaluating its feasibility and efficacy modulating AD’s progression in combination with the naturally derived anti-inflammatory quercetin (Advani & Kumar, 2021; Gonzales et al., 2022). Blood cancers and rheumatoid arthritis drugs with anti-inflammatory properties (from a pool of 80 FDA-approved and clinically tested drugs) were already pinpointed as viable repositioned candidates to halt or reduce AD affectations in a whole-brain transcriptomics machine learning approach (Rodriguez et al., 2021). Here, we have delved into the molecular mechanisms linked to the synergistic, across-brain pathologies’ *in-vivo* impact on neuronal activity and expanded the search for disease-modifying agents to the entire LINCS database (thousands of perturbagens at a variety of time points, doses, and cell lines) (Evangelista et al., 2022; Xie et al., 2022).

Yet, more specialized implementations would also consider disease heterogeneity, detecting sub-trajectories (Iturria-Medina et al., 2020, 2021) over the AD spectrum and obtaining molecular affectation signatures for each of those phenotypes, thus likely increasing effectiveness in RCTs. In the future, clinicians may target the patient’s unique pathological biomarkers with combination therapies and pleiotropic drugs having universal disease modifying outcomes. Computational drug repositioning may facilitate the process of bringing more effective AD therapy into clinical practice.

## Materials and Methods

### Participants

Data was collected under the Translational Biomarkers in Aging and Dementia (TRIAD) cohort (https://triad.tnl-mcgill.com/). The study was approved by the McGill University PET Working Committee and the Douglas Mental Institute Research Ethics Boards and all participants gave written consent. We selected baseline assessments for 47 “cognitively unimpaired” and 16 “Alzheimer’s disease” subjects (Supplementary File 1—table 1) according to clinical and pathophysiological diagnoses. All subjects underwent T1-weighted structural MRI, resting-state fMRI, Aβ (18F-NAV4694)- and tau (18F-MK-6240)-PET scans –see below, (Sanchez-Rodriguez et al., 2023) and the provided references for processing details. The CU individuals were both Aβ and tau-negative while the selected AD subjects presented positive Aβ status (as determined visually by consensus of two neurologists blinded to the diagnosis) and cortical tau involvement (Braak et al., 1995).

### Image processing

#### MRI

Brain structural T1-weighted 3D images were acquired in sagittal plane for all subjects on a 3 T Siemens Magnetom scanner using a standard head coil with 1 mm isotropic resolution, TE = 2.96 ms, TR = 2300 ms, slice thickness = 1 mm, flip angle = 9 deg, FOV = 256 mm, 192 slices per slab. The images were processed following a standard pipeline (Iturria-Medina et al., 2018) including: non-uniformity correction using the N3 algorithm, segmentation into grey matter, white matter and cerebrospinal fluid (CSF) probabilistic maps (SPM12, www.fil.ion.ucl.ac.uk/spm) and standardization of grey matter segmentations to the MNI space (Evans et al., 1994) using the DARTEL tool (Ashburner, 2007). The images were mapped to the Desikian-Killiany-Touriner (DKT) (Klein & Tourville, 2012) atlas for grey matter segmentation. We selected 66 (bilateral) cortical regions (Supplementary File 1—table 3) that do not present PET off-target binding (Vogel et al., 2020).

#### fMRI

The resting-state fMRI acquisition parameters were: Siemens Magnetom Prisma, echo planar imaging, 860 time points, TR = 681 ms, TE = 32.0 ms, flip angle = 50 deg, number of slices = 54, slice thickness = 2.5 mm, spatial resolution = 2.5×2.5×2.5 mm^3^, EPI factor = 88. We applied a minimal processing pipeline (Iturria-Medina et al., 2018) including motion correction, spatial normalization to the MNI space (Evans et al., 1994) and detrending. We then transformed the signals for each voxel to the frequency domain and computed the ratio of the power in the low-frequency range (0.01–0.08 Hz) to that of the entire BOLD frequency range (0–0.25 Hz), i.e., the fractional amplitude of low-frequency fluctuations (fALFF) (Jia et al., 2019; Yang et al., 2018) – a proxy indicator with high sensibility to disease progression. The fALFF values were averaged over all voxels belonging to a brain region to yield a single value per region.

#### Diffusion Weighted MRI (DW-MRI)

Additionally, high angular resolution diffusion imaging (HARDI) data was acquired for N = 128 cognitively unimpaired subjects in the Alzheimer’s Disease Neuroimaging Initiative (ADNI) (adni.loni.usc.edu). The authors obtained approval from the ADNI Data Sharing and Publications Committee for data use and publication, see documents http://adni.loni.usc.edu/wp-content/uploads/how_to_apply/ADNI_Data_Use_Agreement.pdf and http://adni.loni.usc.edu/wp-content/uploads/how_to_apply/ADNI_Manuscript_Citations.pdf, respectively (Iturria-Medina et al., 2018). For each diffusion scan, 46 separate images were acquired, with 5 b0 images (no diffusion sensitization) and 41 diffusion-weighted images (b = 1000 s/mm2). ADNI aligned all raw volumes to the average b0 image, corrected head motion and eddy current distortions. By using a fully automated fiber tractography algorithm (Iturria-Medina et al., 2007) and intravoxel fiber distribution reconstruction (Tournier et al., 2008), we built region-to-region anatomical connection density matrices where each entry, *C_lk_*, reflects the fraction of the region’s surface involved in the axonal connection with respect to the total surface of both regions, *l* and *k*. Finally, we obtained a representative anatomical network by averaging all the subject-specific connectivity matrices (Sanchez-Rodriguez et al., 2021). Additional details are available in a previous publication where the data was processed and utilized (Iturria-Medina et al., 2018).

#### PET

Study participants had Aβ (^18^F-NAV4694) and tau (^18^F-MK-6240) PET imaging in a Siemens high-resolution research tomograph. ^18^F-NAV4694 images were acquired approximately 40-70 min after the intravenous bolus injection of the radiotracer and reconstructed using an ordered subset expectation maximization (OSEM) algorithm on a 4D volume with three frames (3 × 600 s) (Therriault et al., 2021). ^18^F-MK-6240 PET scans of 20 min (4 × 300 s) were acquired at 90-110 min post-injection (Pascoal et al., 2020). Images were corrected for attenuation, motion, decay, dead time and random and scattered coincidences and, consequently, spatially normalized to the MNI space using the linear and nonlinear registration parameters obtained for the participants’ structural T1 images. ^18^F-MK-6240 images were meninges-striped in native space before performing any transformations to minimize the influence of meningeal spillover. Standardized Uptake Value Ratios (SUVR) for the DKT grey matter regions were calculated using the cerebellar grey matter as the reference region (Iturria-Medina et al., 2018).

### Estimating Aβ and tau-induced neuronal activity alterations

The subject-specific pathophysiological brain activity was computationally generated through coupled Wilson-Cowan (WC) modules with regional firings mediated by Aβ plaques, tau tangles and the interaction of Aβ and tau (modeled as the product of their across-brain deposition levels) (Sanchez-Rodriguez et al., 2023). Each brain region was dynamically represented through coupled excitatory and inhibitory neural masses (Daffertshofer & van Wijk, 2011; Gjorgjieva et al., 2016; Meijer et al., 2015; van Nifterick et al., 2022; Wilson & Cowan, 1972). Unspecific local inputs and cortico-cortical connections additionally stimulated the excitatory populations. The integration of all inputs was achieved by means of a sigmoidal activation function. In our model, the region-specific excitatory firing thresholds in these sigmoid functions depend on the regions’ accumulation of each pathological factor, an assumption based on findings suggesting neuronal excitability changes due to Aβ and/or tau deposition and the much larger excitatory prevalence in the cortex (Busche & Hyman, 2020; Maestú et al., 2021; Targa Dias Anastacio et al., 2022; Tok et al., 2022; van Nifterick et al., 2022; Vossel et al., 2017). Simplistically, we wrote the effective excitatory firing parameter of participant *j* at brain region *k* as linear fluctuations from the normal baseline value (θ_0_) due to the considered pathophysiological factors:

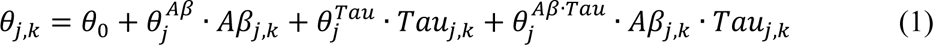

Where 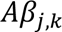 and 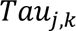 denote the SUVRs normalized to the [0,1] interval –to preserve the dynamical properties of the desired solution–, 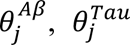 and 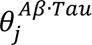 are the brain-wide pathophysiological factor’s influences and each term 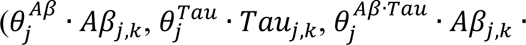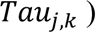 represents the overall factor’s contribution to neuronal activity in subject *j*’s region *k*.

To estimate these pathophysiological contributions, we simulated BOLD signals. The total action potential arriving to the neuronal populations from other local and external populations (Logothetis et al., 2001) underwent metabolic and hemodynamic transformations following Sotero et al. (Sotero & Trujillo-Barreto, 2007, 2008; Valdes-Sosa et al., 2009) to generate the BOLD signal. Then, the parameters maximizing the similarity between the real and simulated individual BOLD signals indicators were obtained via surrogate optimization in MATLAB 2021b (The MathWorks Inc., Natick, MA, USA). The full set of differential equations describing these biophysical transformations and operations is provided in Appendix 1 and (Sanchez-Rodriguez et al., 2023). The equations were solved with an explicit Runge-Kutta (4,5) method, ode45, and a timestep of 0.001s.

Having obtained the likely individual brain-wide influences due to each of the pathological factors (Aβ, tau and Aβ·tau), across-brain mechanistic group differences (AD vs CU) were quantified via the (non-parametric) rank sum test statistics. First, for each subject *j* and brain region, *k*, each pathological factor’s perturbation to neuronal activity in subject *j*’s region *k* was normalized as 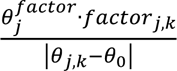. Then, the across-regions vectors resulting from the statistical tests (AD vs CU) quantified the Aβ, tau and Aβ·tau spatial influences on neuronal activity due to AD.

### Neurotypical gene expression profiles

Microarray mRNA expression data from six neurotypical adult brains was downloaded from the Allen Institute (RRID:SCR_007416) website (http://www.brain-map.org). The data was preprocessed by the Allen Institute to reduce the effects of bias due to batch effects (Allen Human Brain Atlas, 2013). For each brain, there were 58,692 probes representing 20,267 unique genes. For genes with multiple probes, Gaussian kernel regression (Gryglewski et al., 2018) was applied to predict the mRNA intensity in each of the 3702 samples in MNI space (Evans et al., 1994) using leave-one-out cross-validation. The probe with the highest prediction accuracy was chosen as the representative probe for that gene. Gaussian kernel regression using mRNA values of proximal regions also served to predict the gene expression for grey matter voxels without mRNA expression intensity. Thus, the whole-brain gene expression data was obtained for the selected 20,267 probes/genes. Probes/genes described as “uncharacterized”, “similar to hypothetical protein”, “pseudogene” were dropped, leaving 19,469. Finally, we calculated average gene expression values for each region in the brain parcellation (Adewale et al., 2021).

### Molecular associates of the Aβ, tau and Aβ·tau spatial alterations to neuronal activity

We aimed to determine the genes with whole-brain expressions predicting the Aβ, tau and Aβ·tau effects on neuronal activity. For each pathological factor, we evaluated monotonic relationships between the corresponding neuronal activity spatial alterations patterns and the regional gene expression values by computing Spearman correlations. We estimated 99% Spearman’s rho confidence intervals with 100,000 bootstrapping resamples and retained the genes which confidence limits did not include zero (significant correlation). The resulting sets of genes were termed Aβ, tau and Aβ·tau molecular associates, respectively.

### Statistical analyses

We performed functional pathways enrichment analyses on Metascape (Zhou et al., 2019), a web-based portal that integrates various independent biological databases (KEGG Pathway, GO Biological Processes, Reactome Gene Sets, Canonical Pathways, CORUM, WikiPathways, PANTHER Pathway, DisGeNET), using default specifications. Metascape identifies ontology terms that are significantly over-represented in the input gene lists through hypergeometric tests and the Benjamini-Hochberg *p*-value correction algorithm (*q < 0.05*). To avoid redundancy from the reporting of multiple ontologies, Kappa similarities among all pairs of enriched terms are computed. Then, the similarity matrix is hierarchically clustered, and a 0.3 threshold is applied. The most significant (lowest p-value) term within each cluster is chosen to represent the cluster (Zhou et al., 2019). Cell type enrichment was performed with the Expression Weighted Celltype Enrichment toolbox (Skene & Grant, 2016). The probability of enrichment is determined as the percentage of 100,000 random gene lists in a background set with lower average expression in each cell type than in our gene lists. The background gene set is comprised of all genes with orthologs between human and mice and its single-cell transcriptome data were sampled from the mice somatosensory cortex and hippocampus CA1 (Skene & Grant, 2016). Drug repurposing alternatives were investigated on the webserver SigCom LINCS (Evangelista et al., 2022; Xie et al., 2022). This search engine uses a database of ranked gene lists for drug-induced gene expression changes. Similarity and statistical measures (p-values, Benjamini-Hochberg corrected, *q < 0.05*) are computed using the Mann–Whitney U test: the average rank of the user-provided gene set in each chemical perturbation’s gene list is compared to the average rank of a randomly selected gene set (Evangelista et al., 2022; Xie et al., 2022).

## Supporting information

Supplementary file 2: A&#946, tau and A&#946*tau functional signatures gene lists.

Supplementary file 3: Gene ontology enrichment analysis of the combined A&#946, tau and A&#946*tau signature gene sets (using Metascape).

Supplementary file 4: Gene ontology enrichment analysis of the A&#946 signature gene set (using Metascape).

Supplementary file 5: Gene ontology enrichment analysis of the tau signature gene set (using Metascape).

Supplementary file 6: Gene ontology enrichment analysis of the A&#946*tau signature gene set (using Metascape).

Supplementary file 7: Gene-disease associations enrichment analysis of the A&#946 signature gene set (DisGeNET, using Metascape).

Supplementary file 8: Gene-disease associations enrichment analysis of the tau signature gene set (DisGeNET, using Metascape).

Supplementary file 9: Gene-disease associations enrichment analysis of the A&#946*tau signature gene set (DisGeNET, using Metascape).

Supplementary file 10: Top chemical perturbations that up-regulate the combined A&#94, tau and A&#946*tau signature gene sets (using SigCom LINCS).

Supplementary file 11: Top chemical perturbations that down-regulate the combined A&#, tau and A&#946*tau signature gene sets (using SigCom LINCS).

Supplementary file 12: Drug indications and BBB permeability of the FDA-approved top chemical perturbations that up and down-regulate the gene sets

Supplementary file 13: Top chemical perturbations that up-regulate the A&#946 signature gene set (using SigCom LINCS).

Supplementary file 14: Top chemical perturbations that down-regulate the A&#946 signature gene set (using SigCom LINCS).

Supplementary file 15: Top chemical perturbations that up-regulate the tau signature gene set (using SigCom LINCS).

Supplementary file 16: Top chemical perturbations that down-regulate the tau signature gene set (using SigCom LINCS).

Supplementary file 17: Top chemical perturbations that up-regulate the A&#946*tau signature gene set (using SigCom LINCS).

Supplementary file 18: Top chemical perturbations that down-regulate the A&#946*tau signature gene set (using SigCom LINCS).

Supplementary file 1: Supplementary Tables 1-3, Supplementary Figure 1 and Supplementary Pseudocode 1.

Appendix 1: Personalized AD neuronal activity model

## Data availability

The data that support the findings of this study are available by submitting a data share request via https://triad.tnl-mcgill.com/contact-us/. All the data collected under the TRIAD cohort is governed by the policies set by the Research Ethics Board Office of the McGill University, Montreal and the Douglas Research Center, Verdun.

## Code availability

The code utilized in this article for the neuronal activity simulations and quantification of the pathological effects will be freely available with publication at the *Neuroinformatics for Personalized Medicine* lab’s website (NeuroPM, https://www.neuropm-lab.com/publication-codes.html). Supplementary File 1—pseudocode 1 contains the algorithm’s overview.

## Acknowledgments

We thank Lisa Münter for valuable feedback on the manuscript draft. LSR was partially supported by funding from the Fonds de recherche du Québec – Santé. This project was undertaken thanks in part to the following funding awarded to YIM: the Canada Research Chair tier-2, the CIHR Project Grant 2020, and the Weston Family Foundation’s AD Rapid Response 2018 and Transformational Research in AD 2020. In addition, we used the computational infrastructure of the McConnell Brain Imaging Center at the Montreal Neurological Institute, supported in part by the Brain Canada Foundation, through the Canada Brain Research Fund, with the financial support of Health Canada and sponsors. PRN, GB, JT, JF, SS, NR, CT and JS are supported by the Canadian Institutes of Health Research (CIHR) [MOP-11-51-31; RFN 152985, 159815, 162303], Canadian Consortium of Neurodegeneration and Aging (CCNA; MOP-11-51-31 -team 1), Weston Brain Institute, the Alzheimer’s Association [NIRG-12-92090, NIRP-12-259245], Brain Canada Foundation (CFI Project 34874; 33397), the Fonds de Recherche du Québec – Santé (FRQS; Chercheur Boursier, 2020-VICO-279314) and the Colin J. Adair Charitable Foundation.

## List of Supplementary Files

**Supplementary file 1**

**Supplementary Tables 1-3, Supplementary Figure 1 and Supplementary Pseudocode 1.**

**Supplementary file 2**

**Aβ, tau and Aβ·tau functional signatures gene lists.**

**Supplementary file 3**

**Gene ontology enrichment analysis of the combined Aβ, tau and Aβ·tau signature gene sets (using Metascape).**

**Supplementary file 4**

**Gene ontology enrichment analysis of the Aβ signature gene set (using Metascape).**

**Supplementary file 5**

**Gene ontology enrichment analysis of the tau signature gene set (using Metascape).**

**Supplementary file 6**

**Gene ontology enrichment analysis of the Aβ·tau signature gene set (using Metascape).**

**Supplementary file 7**

**Gene-disease associations enrichment analysis of the Aβ signature gene set (DisGeNET, using Metascape).**

**Supplementary file 8**

**Gene-disease associations enrichment analysis of the tau signature gene set (DisGeNET, using Metascape).**

**Supplementary file 9**

**Gene-disease associations enrichment analysis of the Aβ·tau signature gene set (DisGeNET, using Metascape).**

**Supplementary file 10**

**Top chemical perturbations that up-regulate the combined Aβ, tau and Aβ·tau signature gene sets (using SigCom LINCS).**

**Supplementary file 11**

**Top chemical perturbations that down-regulate the combined Aβ, tau and Aβ·tau signature gene sets (using SigCom LINCS).**

**Supplementary file 12**

**Drug indications and blood-brain barrier permeability of the FDA-approved top chemical perturbations that up and down-regulate the combined Aβ, tau and Aβ·tau signature gene sets (using SigCom LINCS, PubChem and B3DB).**

**Supplementary file 13**

**Top chemical perturbations that up-regulate the Aβ signature gene set (using SigCom LINCS).**

**Supplementary file 14**

**Top chemical perturbations that down-regulate the Aβ signature gene set (using SigCom LINCS).**

**Supplementary file 15**

**Top chemical perturbations that up-regulate the tau signature gene set (using SigCom LINCS).**

**Supplementary file 16**

**Top chemical perturbations that down-regulate the tau signature gene set (using SigCom LINCS).**

**Supplementary file 17**

**Top chemical perturbations that up-regulate the Aβ·tau signature gene set (using SigCom LINCS).**

**Supplementary file 18**

**Top chemical perturbations that down-regulate the Aβ·tau signature gene set (using SigCom LINCS).**

