## Supplementary file 1: Supplementary Tables 1-3, Supplementary Figure 1 and Supplementary Pseudocode 1. for "Transcriptomic signatures of Aβ- and tau-induced neuronal dysfunction reveal inflammatory processes at the core of Alzheimer’s disease pathophysiology"

Sanchez-Rodriguez et al.

**Supplementary file 1—table 1.** Demographics of the samples

| | CU | AD | p |
|---|---|---|---|
| Number of individuals (N) | 47 | 16 | - |
| Age (yrs), mean (s.d.) | 68.84 (8.36) | 69.45 (9.10) | 0.801 |
| Female, N (%) | 36 (76.6) | 8 (50.0) | 0.061 |
| Education (yrs), mean (s.d.) | 15.64 (3.59) | 14.31 (3.38) | 0.210 |
| *APOE* ε4 carriers, N (%) | 11 (23.4) | 7 (43.8) | 0.198 |
| MMSE, mean (s.d.) | 29.33 (0.88) | 20.25 (7.20) | < 0.001 |
| Aβ+, N (%) | 0 (0) | 16 (100.0) | < 0.001 |

The reported p-values are for comparisons to cognitively unimpaired (CU) subjects. P-values for age, education and MMSE indicate values assessed with two-sided independent-samples t-tests. For the resting variables (sex, *APOE* ε4 status and Aβ+), Fischer exact tests were performed. CU cognitively unimpaired; AD Alzheimer's disease; *APOE* ε4, apolipoprotein epsilon 4; MMSE, Mini-Mental State examination.

**Supplementary file 1—table 2.** Genes with most appearances (%) in the top statistically significant biological pathways identified for the Aβ+tau → neuronal-activity gene list.

| Gene | % | Gene | % | Gene | % | Gene | % | Gene | % |
|---|---|---|---|---|---|---|---|---|---|
| *RIPK2* | 45.7 | *CLEC7A* | 21.7 | *TICAM2* | 17.2 | *SLAMF6* | 14.5 | *C5AR1* | 12.7 |
| *SYK* | 45.2 | *TYROBP* | 21.7 | *RPH3AL* | 17.2 | *TLR6* | 14.5 | *OSR1* | 12.2 |
| *ANXA1* | 41.2 | *SNCA* | 21.3 | *MAPK1* | 17.2 | *VAMP8* | 14.5 | *HIPK2* | 12.2 |
| *IL12B* | 38.0 | *NAGLU* | 21.3 | *ITPKB* | 17.2 | *SRF* | 14.5 | *CD84* | 12.2 |
| *CCL19* | 36.2 | *CD1D* | 21.3 | *HLA-DRA* | 17.2 | *PTCH1* | 14.5 | *FCN3* | 12.2 |
| *HLA-DRB1* | 35.7 | *ZBTB16* | 20.8 | *CD38* | 17.2 | *PLA2G5* | 14.5 | *NCK2* | 12.2 |
| *CCL5* | 33.0 | *VAV1* | 20.8 | *GAL* | 16.7 | *HLA-DPB1* | 14.5 | *TNFRSF4* | 12.2 |
| *IHH* | 32.6 | *B2M* | 20.4 | *BMP7* | 16.7 | *CCR1* | 14.5 | *TLR1* | 12.2 |
| *PTPRC* | 31.7 | *WNT10B* | 19.9 | *SPHK2* | 16.3 | *TBX3* | 14.0 | *SKI* | 12.2 |
| *CD74* | 31.7 | *IL2RG* | 19.9 | *PLCB1* | 16.3 | *OPRK1* | 14.0 | *PIK3R1* | 12.2 |
| *IL1B* | 31.2 | *GRP* | 19.9 | *CD160* | 16.3 | *CRHBP* | 14.0 | *HLA-DPA1* | 12.2 |
| *IL18* | 30.8 | *F2* | 19.9 | *EBI3* | 16.3 | *AIF1* | 14.0 | *MTOR* | 12.2 |
| *TGFB1* | 29.9 | *EPO* | 19.9 | *SOX11* | 16.3 | *FCRL3* | 13.6 | *CPLX1* | 11.8 |
| *F2RL1* | 29.9 | *NF1* | 19.5 | *PIK3CG* | 16.3 | *CYP26B1* | 13.6 | *PAX2* | 11.8 |
| *BMP4* | 29.9 | *KITLG* | 19.5 | *NPY* | 16.3 | *BAIAP3* | 13.6 | *GNAI2* | 11.8 |
| *PYCARD* | 29.4 | *FYN* | 19.5 | *ITGAM* | 16.3 | *SOD1* | 13.6 | *APLNR* | 11.8 |
| *PLCG2* | 28.5 | *GHRL* | 19.0 | *IL1A* | 16.3 | *PTGER4* | 13.6 | *SYT2* | 11.3 |
| *NCKAP1L* | 26.7 | *GPR68* | 19.0 | *CD36* | 16.3 | *NTSR1* | 13.6 | *CADPS2* | 11.3 |
| *EDN1* | 26.2 | *CCL2* | 19.0 | *CBFB* | 16.3 | *CRH* | 13.6 | *SYT12* | 11.3 |
| *CD80* | 26.2 | *PTAFR* | 19.0 | *OPRM1* | 15.8 | *BBS4* | 13.6 | *RPH3A* | 11.3 |
| *CD86* | 25.8 | *CFTR* | 19.0 | *C3* | 15.8 | *TLR7* | 13.1 | *FGL2* | 11.3 |
| *HLA-G* | 25.3 | *RBP4* | 18.6 | *PELI1* | 15.4 | *MBL2* | 13.1 | *PDE5A* | 11.3 |
| *HAVCR2* | 24.9 | *INS* | 18.6 | *CITED2* | 15.4 | *ADCYAP1* | 13.1 | *EOMES* | 11.3 |
| *SOX4* | 24.9 | *LILRB2* | 18.1 | *KIAA0748* | 15.4 | *TNIP2* | 12.7 | *LAPTM5* | 11.3 |
| *WNT3A* | 24.4 | *STXBP1* | 18.1 | *RAG1* | 15.4 | *CPLX2* | 12.7 | *SPINK1* | 11.3 |
| *HLA-E* | 24.4 | *GPR183* | 18.1 | *LRP5* | 15.4 | *NRP1* | 12.7 | *PSMD9* | 11.3 |
| *WNT7A* | 24.0 | *IRF4* | 17.6 | *GATA2* | 15.4 | *KCNQ1* | 12.7 | *NOS2* | 11.3 |
| *KIT* | 24.0 | *HLA-F* | 17.6 | *ACTL6A* | 15.4 | *IL12A* | 12.7 | *CHGA* | 11.3 |
| *CARD11* | 23.1 | *FGB* | 17.6 | *STXBP3* | 14.9 | *CD70* | 12.7 | *CEBPA* | 11.3 |

**Supplementary file 1—table 3.** Brain regions in the considered parcellation.

| | | |
|---|---|---|
| Hippocampus | Lateral Orbitofrontal | Posterior Cingulate |
| Entorhinal | Medial Orbitofrontal | Precentral |
| Amygdala | Lingual | Precuneus |
| Caudal Anterior Cingulate | Middle Temporal | Rostral Anterior Cingulate |
| Caudal Middle Frontal | Parahippocampal | Rostral Middle Frontal |
| Cuneus | Paracentral | Superior Frontal |
| Fusiform | Pars Opercularis | Superior Parietal |
| Inferior Parietal | Pars Orbitalis | Superior Temporal |
| Inferior Temporal | Pars Triangularis | Supramarginal |
| Isthmus Cingulate | Pericalcarine | Transverse Temporal |
| Lateral Occipital | Postcentral | Insula |

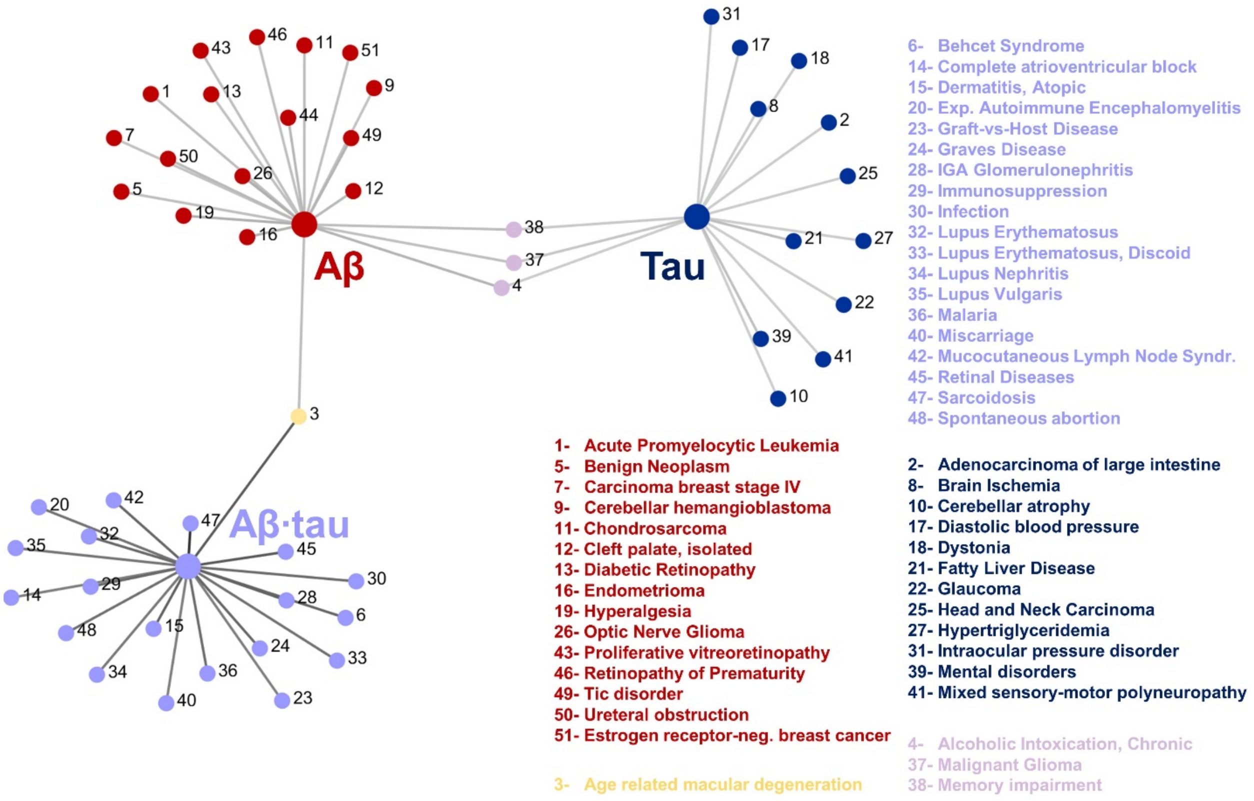


**Supplementary file 1—figure 1. AD associated diseasome.** To visually represent the associations between the AD Aβ, tau and Aβ·tau molecular associates and other disease-characteristic molecular pathways, the grayscale color of a link in the network plot is inversely proportional to the q-value of the enrichment statistical test, i.e., darker edges reflect increased statistical significance that the dysfunctional disease pathways are overrepresented in the corresponding gene set. Only significantly enriched terms are shown (hypergeometric tests, $q < 0.05$, Benjamini-Hochberg corrected), to a maximum of 20 terms. Top disease pathways that were shared by the functional genetic signatures connect the Aβ, tau and Aβ·tau clusters, being represented with additional colors. The network plot was annotated on the right with the names of the associated pathologies for increased readability.

**Supplementary file 1—pseudocode 1.**

```
// program to calculate the personalized combined neuronal activity influences by Aβ, tau and
// Aβ · Tau

{       // definitions
Define surrogate optimization parameters
Load the subjects Aβ, tau and fALFF (rs-fMRI) and anatomical connectivity matrix
Define the neuronal activity influence model (Eq. 1)
Define neural mass model and transformations to simulate the resting-state BOLD signal
Define the objective function (minimizes distance between real and simulated BOLD)
}

{       // optimization
FOR i = 1 TO 20       // different random optimization evaluation trials
        Perform surrogate optimization until the algorithm converges
                // At each iteration:
                // simulate the BOLD signal,
                // calculate similarity with the subjects real signal,
                // retain the best evaluation thus far
                // (performed by Matlab's surrogateopt.m)
        Save the optimized neuronal activity affectation parameters and optimization outputs
ENDFOR
}

{       // post-processing
Retain the optimization outcome with the lowest overall cost
Reconstruct hidden quantities of interest, e.g., Aβ, tau and Aβ·tau subject-specific spatial effects
on neuronal activity
}
```
