## Appendix 1: Personalized AD neuronal activity model for "Transcriptomic signatures of Aβ- and tau-induced neuronal dysfunction reveal inflammatory processes at the core of Alzheimer’s disease pathophysiology"

For each participant, the BOLD signal is generated through coupled differential equations. Firstly, the excitatory and inhibitory firing rates (Daffertshofer & van Wijk, 2011; Gjorgjieva et al., 2016; Wilson & Cowan, 1972) in neural mass $k$, $E_k(t)$ and $I_k(t)$, are obtained from:

$$\dot{E}_k = \frac{1}{\tau_E}\left[-E_k + S\left(x_{E,k}\right)\right]$$

$$\dot{I}_k = \frac{1}{\tau_I}\left[-I_k + S\left(x_{I,k}\right)\right]$$

$$x_{E,k} = C_{EE}E_k - C_{IE}I_k + P + \frac{\eta}{N}\sum_{l=1,l\neq k}^{N} \mathrm{C}_{lk}E_l$$

$$x_{I,k} = C_{EI}E_k - C_{II}I_k$$

with the sigmoidal activation functions $S_I\left(x_{I,k}\right) = \frac{1}{1+\exp\left[-a_I\left(x_{I,k}-\theta_I\right)\right]} - \frac{1}{1+\exp\left[a_I\theta_I\right]}$ and $S_E\left(x_{E,k}\right) = \frac{1}{1+\exp\left[-a_E\left(x_{E,k}-\theta_{E,k}\right)\right]} - \frac{1}{1+\exp\left[a_E\theta_{E,k}\right]}$. Additionally, each excitatory sigmoidal firing threshold depends on the local amyloid-beta ($A\beta$) and tau loads: $\theta_{E,k} = \theta_0 + \theta_E^{A\beta} \cdot A\beta_k + \theta_E^{Tau} \cdot Tau_k + \theta_E^{A\beta\cdot Tau} \cdot A\beta_k \cdot Tau_k$. This equation is as (1) in the main text, highlighting that the model assumes perturbations to the excitatory parameter by the pathogens. It must be understood that the $A\beta$ and tau accumulations are subject-specific.

The BOLD signal relates to the action potential arriving at the neuronal populations (Logothetis et al., 2001; Sotero & Trujillo-Barreto, 2008; Valdes-Sosa et al., 2009). All quantities are normalized to baseline values: $\xi_{E,k} = \frac{S_{E,k}}{{S_{E,k}}^0}$ and $\xi_{I,k} = \frac{S_{I,k}}{{S_{I,k}}^0}$, where the superscript denotes values at rest.

Changes in glucose consumption ($g_{E,k}$ and $g_{I,k}$) are linked to the excitatory and inhibitory neuronal inputs in region $k$. The glucose variables transform into metabolic rates of oxygen for excitatory ($m_{E,k}$) and inhibitory ($m_{I,k}$) activities, and total oxygen consumption ($m_k$):

$$\dot{g}_{E,k} = z_{E,k}$$

$$\dot{z}_{E,k} = \frac{-2}{\varkappa_E} z_{E,k} - \frac{1}{{\varkappa_E}^2}\left(g_{E,k} - 1\right) + \frac{h_E}{\varkappa_E}\left(\xi_{E,k} - 1\right)$$

$$\dot{g}_{I,k} = z_{I,k}$$

$$\dot{z}_{I,k} = \frac{-2}{\varkappa_I} z_{I,k} - \frac{1}{{\varkappa_I}^2}\left(g_{I,k} - 1\right) + \frac{h_I}{\varkappa_I}\left(\xi_{I,k} - 1\right)$$

$$m_{E,k}(t) = \frac{2 - x(t)}{2 - x_0} g_{E,k}(t)$$

$$m_{I,k}(t) = g_{I,k}(t)$$

$$m_k(t) = \frac{\gamma m_{E,k}(t) + m_{I,k}(t)}{\gamma + 1}$$

$$x(t) = \frac{1}{1 + exp\left[c\left(d - g_{E,k}(t)\right)\right]}$$

Cerebral blood flow ($f_k$) is modeled as follows (Friston et al., 2000), assuming that CBF is coupled to excitatory activity:

$$\dot{f}_k = y_k$$

$$\dot{y}_k = \frac{-2}{\varkappa_f} y_k - \frac{1}{{\varkappa_f}^2}(f_k - 1) + \mu\left(\xi_{E,k} - 1\right)$$

The outputs of the metabolic and vascular models are converted to normalized cerebral blood volume ($b_k$) and deoxy-hemoglobin ($q_k$) content through the Balloon model (Buxton et al., 1998):

$$\dot{b}_k = \frac{1}{\varkappa_0}(f_k - f_{out})$$

$$\dot{q}_k = \frac{1}{\varkappa_0}\left(m_k - f_{out}\frac{q_k}{b_k}\right)$$

$$f_{out} = b_k^{\frac{1}{\zeta}}$$

The BOLD signal is finally obtained by using the following linear observation equation:

$$BOLD_k(t) = V_0\big(a_1(1 - q_k) - a_2(1 - b_k)\big)$$

where $a_1 = 4.3Y_0E_0 \cdot TE + \varepsilon r_0 E_0 \cdot TE$ and $a_2 = \varepsilon r_0 E_0 \cdot TE + \varepsilon - 1$ are parameters that depend on the experimental conditions (field strength, $TE$) (Archila-Meléndez et al., 2020; Deco et al., 2018; Obata et al., 2004; Simon & Buxton, 2015).

**Appendix 1—table 1.** Dynamical model parameters

| Parameter | Definition | Value | Ref. |
|---|---|---|---|
| $\tau_I$ | Time-constant controlling the decay of inhibitory activity after stimulation | 0.02 s | (Abeysuriya et al., 2018) |
| $\tau_E$ | Time-constant controlling the decay of excitatory activity after stimulation | 0.01 s | (Abeysuriya et al., 2018) |
| $C_{II}$ | Local inhibitory-inhibitory connection strength | 1.2 | (Gjorgjieva et al., 2016; Meijer et al., 2015; Wilson & Cowan, 1972) |
| $C_{EI}$ | Local excitatory-inhibitory connection strength | 6 | (Gjorgjieva et al., 2016; Meijer et al., 2015; Wilson & Cowan, 1972) |
| $C_{EE}$ | Local excitatory-excitatory connection strength | 6.4 | (Gjorgjieva et al., 2016; Meijer et al., 2015; Wilson & Cowan, 1972) |
| $C_{IE}$ | Local inhibitory-excitatory connection strength | 4.8 | (Gjorgjieva et al., 2016; Meijer et al., 2015; Wilson & Cowan, 1972) |
| $P$ | Average constant external input received by the excitatory population | 0.65<br>(set to produce plausible simulated electrophysiological and BOLD signals) | (Gjorgjieva et al., 2016; Meijer et al., 2015; Wilson & Cowan, 1972) |
| $a_I$ | Maximum slope of the inhibitory sigmoidal activation function | 1 | (Abeysuriya et al., 2018) |
| $a_E$ | Maximum slope of the excitatory sigmoidal activation function | 1 | (Abeysuriya et al., 2018) |
| $\theta_I$ | Position of the inhibitory sigmoidal firing function' threshold for activation | 4 | (Gjorgjieva et al., 2016; Meijer et al., 2015; Wilson & Cowan, 1972) |
| $\theta_E$ | Position of the excitatory sigmoidal firing function' threshold for activation | Variable in [2.75,2.85] depending on the regional pathological loads<br><br>2.8 (in normal baseline conditions) | (Abeysuriya et al., 2018; Daffertshofer & van Wijk, 2011; Gjorgjieva et al., 2016; Meijer et al., 2015; Wilson & Cowan, 1972) |
| $\eta$ | Global coupling strength scaling the anatomical connectivity matrix $C_{lk}$ | 2<br>(set to produce plausible simulated electrophysiological and BOLD signals) | (Abeysuriya et al., 2018; Daffertshofer & van Wijk, 2011; Gjorgjieva et al., 2016; Meijer et al., 2015; Wilson & Cowan, 1972) |
| $N$ | Number of brain regions of interest | 66 | (Klein & Tourville, 2012) |

| | | | |
|---|---|---|---|
| $h_E$ | Efficacy of glucose consumption response to excitation | 1 | (Sotero et al., 2009; Sotero & Trujillo-Barreto, 2007, 2008; Valdes-Sosa et al., 2009) |
| $h_I$ | Efficacy of glucose consumption response to inhibition | 1 | (Sotero et al., 2009; Sotero & Trujillo-Barreto, 2007, 2008; Valdes-Sosa et al., 2009) |
| $\varkappa_E$ | Time-constant of the excitatory glucose consumption impulse response. | 1 s | (Sotero et al., 2009; Sotero & Trujillo-Barreto, 2007, 2008; Valdes-Sosa et al., 2009) |
| $\varkappa_I$ | Time-constant of the inhibitory glucose consumption impulse response. | 1 s | (Sotero et al., 2009; Sotero & Trujillo-Barreto, 2007, 2008; Valdes-Sosa et al., 2009) |
| $c$ | Steepness of the sigmoid function $x$ | 2.5 | (Sotero et al., 2009; Sotero & Trujillo-Barreto, 2007, 2008; Valdes-Sosa et al., 2009) |
| $d$ | Position of the threshold of the sigmoid function $x$ | 1.6 | (Sotero et al., 2009; Sotero & Trujillo-Barreto, 2007, 2008; Valdes-Sosa et al., 2009) |
| $\gamma$ | Baseline ratio of excitatory to inhibitory synaptic activity in the voxel | 5 | (Sotero et al., 2009; Sotero & Trujillo-Barreto, 2007, 2008; Valdes-Sosa et al., 2009) |
| $x_0$ | Fraction of glucose following the glycogenolitic pathway at rest | $\frac{1}{1+exp\left[c\left(d-1(t)\right)\right]}$ | (Sotero et al., 2009; Sotero & Trujillo-Barreto, 2007, 2008; Valdes-Sosa et al., 2009) |
| $\mu$ | Efficacy of blood flow response to excitation | 0.8 | (Sotero et al., 2009; Sotero & Trujillo-Barreto, 2007, 2008; Valdes-Sosa et al., 2009) |
| $\varkappa_f$ | Time constant for CBF response | 1.7 | (Sotero et al., 2009; Sotero & Trujillo-Barreto, 2007, 2008; Valdes-Sosa et al., 2009) |
| $\varkappa_0$ | Transit time through the balloon | 1 | (Sotero et al., 2009; Sotero & Trujillo-Barreto, 2007, 2008; Valdes-Sosa et al., 2009) |
| $\zeta$ | Coefficient of the steady state flow-volume relationship | 0.4 | (Sotero et al., 2009; Sotero & Trujillo-Barreto, 2007, 2008; Valdes-Sosa et al., 2009) |
| $V_0$ | Baseline blood volume | 0.03 | (Sotero et al., 2009; Sotero & Trujillo-Barreto, 2007, 2008; Valdes-Sosa et al., 2009) |
| $\Upsilon_0$ | frequency offset of a fully deoxygenated blood vessel at 3 T | 80.6 $s^{-1}$ (at 3 T) | (Archila-Meléndez et al., 2020; Obata et al., 2004; Simon & Buxton, 2015) |
| $r_0$ | Slope defining the dependence of the R2* relaxation rate on blood oxygenation | 178 $s^{-1}$ (at 3 T) | (Archila-Meléndez et al., 2020; Obata et al., 2004; Simon & Buxton, 2015) |

| | | | |
|---|---|---|---|
| $E_0$ | Baseline oxygen extraction fraction | 0.4 | (Archila-Meléndez et al., 2020; Obata et al., 2004; Simon & Buxton, 2015) |
| $\varepsilon$ | Intrinsic ratio of blood to tissue signals at rest | 0.24 | (Archila-Meléndez et al., 2020; Obata et al., 2004; Simon & Buxton, 2015) |
| $TE$ | Echo time | 32.0 ms | https://triad.tnl-mcgill.com/ |
